# Hematopoietic Tet2 inactivation enhances the response to checkpoint blockade immunotherapy

**DOI:** 10.1101/2024.09.09.612140

**Authors:** Robert J. Vanner, Suraj Bansal, Marco M. Buttigeig, Andy G.X. Zeng, Vincent Rondeau, Darryl Y. Chan, Michelle Chan-Seng-Yue, Liqing Jin, Jessica McLeod, Elisa Donato, Patrick Stelmach, Caitlyn Vlasschaert, Yitong Yang, Aarushi Gupta, Sofia Genta, Enrique Sanz Garcia, Liran Shlush, Mauricio Ribeiro, Marcus O. Butler, Sagi Abelson, Mark Minden, Steven M. Chan, Michael J. Rauh, Andreas Trumpp, John E Dick

## Abstract

Somatic mutations inactivating *TET2* are among the most common drivers of clonal hematopoiesis (CH). While TET2 inactivation is associated with monocyte-derived inflammation and improved chimeric antigen-receptor-T cell function, its impact on immunotherapy response is unknown. In our mouse model, hematopoietic *Tet2* mutation enhanced immune checkpoint blockade (ICB) response. Enhanced ICB response with *Tet2* mutation required phagocytes, CD4 and CD8 T cells. Mechanistically, in *Tet2*-mutant tumor-infiltrating leukocytes (TILs), ICB preferentially induced anti-tumor states and restricted cell states linked to tumor progression. *Tet2*-mutant monocytes activated costimulatory programs, while *Tet2*-mutant T cells showed enhanced T cell memory signatures, lesser exhaustion and decreased regulatory phenotype. Our murine data was clinically relevant, since we found that melanomas from patients with *TET2* driver mutation-CH (TET2-CH) showed enhanced immune infiltration, T cell activation, and T cell memory programs. In melanoma patients treated with ICB, TET2-CH was associated with 6-fold greater odds of clinical benefit. Collectively, our data establishes that hematopoietic Tet2 inactivation primes leukocytes for anti-tumor states associated with immunotherapy response and provides a potential biomarker for personalized therapy.

## INTRODUCTION

Somatic mutations inactivating the epigenetic regulator Tet methylcytosine dioxygenase 2 (*TET2*) in hematopoietic stem cells (HSC) are one of the most common drivers of clonal hematopoiesis (CH) in patients with blood cancers and solid tumors^1^. While CH-mutant leukocytes infiltrate tumors, and are potentially enriched within TILs^2^, little is known about how this influences tumor biology and cancer outcomes. *TET2*-mutant CH (TET2-CH) is linked to accelerated cardiovascular, lung, liver, and renal disease, with animal models demonstrating enhanced inflammation from *Tet2*-mutant monocytes/macrophages as a potential mechanism^3^. Intriguingly, myeloid-specific *Tet2*-knockout curbed melanoma growth in a mouse model, where *Tet2*-deficiency attenuated the immunosuppressive properties of tumor-infiltrating leukocytes (TILs)^4^. *Tet2*-knockout also improved adoptive T cell therapy response in syngeneic and xenograft tumors^5,6^. Strikingly, *TET2* inactivation by viral integration in a CAR-T cell-treated patient led to a complete response with concomitant *TET2*-mutant CAR-T clonal dominance^6^. Since TET2 mutation activates innate and adaptive immunity, and TET2-CH has been associated with gene-expression programs predictive of immunotherapy response^7^, we sought to define the impact of *Tet2* inactivation on immune checkpoint blockade response (ICB).

ICB has revolutionized cancer therapy, where some patients achieve durable disease control, however most either do not respond or eventually relapse^8^. While somatic mutations are known to influence immune effector function, how CH impacts immunotherapy response is unclear. Despite *in vitro* and animal model evidence for enhanced anti-tumor immunity from *Dnmt3a*^9^ or *Tet2*^5^ mutant leukocytes, CH was retrospectively associated with worse overall survival in solid tumor patients treated with ICB^10^. The strong association between CH and prior cancer therapy may confound these analyses since patients with CH are enriched for individuals exposed to more treatment and thus with more advanced disease. Also, cancer-free individuals with CH have worse overall survival largely due to cardiovascular disease, so the prognostic association in cancer patients may not be entirely linked to altered tumor biology. Colorectal cancer patients uniquely showed a positive association between CH and survival post ICB^10^. Patients with colorectal cancer and CH also had improved survival in the FIRE-3 trial of chemotherapy and bevacizumab or cetuximab for newly diagnosed metastatic colorectal cancer, which suggests that CH-associated outcomes are context-specific^11^.

Most studies of CH in oncology patients have considered all driver mutations as a single entity, even though drivers differ in their impact on hematopoietic cell function and association with outcome^3^. Here we use an integrated approach combining syngeneic tumor models with hematopoietic Tet2 inactivation and primary clinical data from ICB-treated patients to assess how TET2-CH modulates patient outcomes and leukocyte biology with ICB.

## RESULTS AND DISCUSSION

### Tet2-CH enhances anti-PD-1 immunotherapy response

We tested the effect of *Tet2*-mutant hematopoiesis on immunotherapy response using a syngeneic mouse colorectal cancer model MC38. We selected MC38 as it does not require foreign neo-antigens for ICB response and CH has been associated with immunotherapy benefit in colorectal cancer^10^. Lethally-irradiated C57Bl6/J mice were rescued with unfractionated bone marrow cells from wild type (Tet2^wt^) or *Tet2* mutant mice. After Tet2^wt^ or *Tet2*-mutant hematopoietic reconstitution, respective recipients were subcutaneously implanted with MC38 cells and tumour growth was monitored without intervention, or in mice treated with isotype control antibody or Programmed cell death protein 1 (PD-1) blocking ICB (Figure 1A). In CD45.1^+^ B6 recipients rescued with CD45.2^+^ Tet2^wt^ or Tet2-heterozygous null (Tet2^het^) bone marrow prior to tumor implantation, TILs were almost universally bone-marrow derived, in keeping with prior reports (Supplementary Figure 1A-1B) ^12^. Without treatment, there was no difference between tumor growth kinetics in Tet2^wt^ or Tet2^het^ mice (Supplementary Figure 1C).

**Figure 1.**
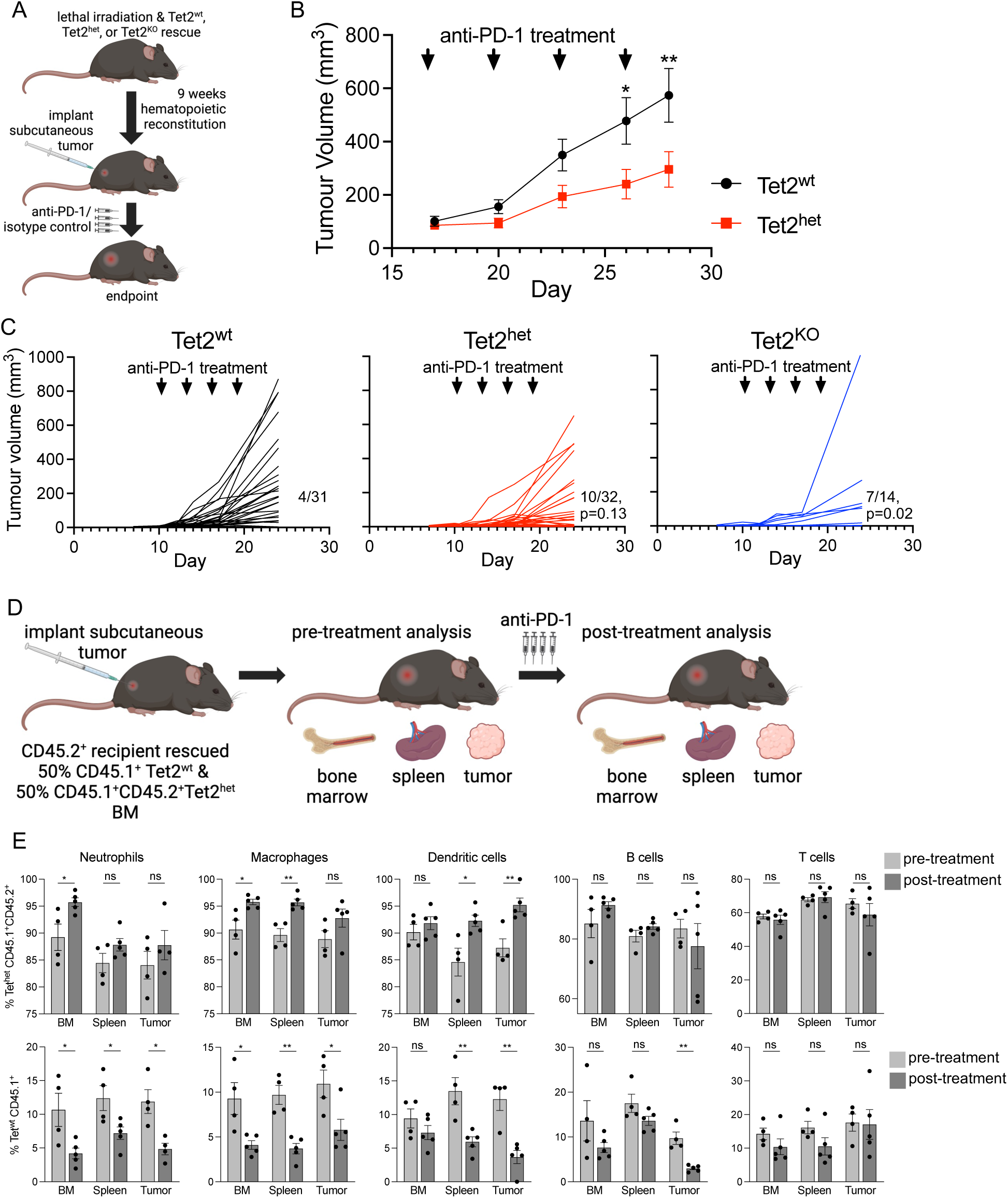
Myeloid-biased *Tet2*-mutant hematopoiesis improves PD-1 Checkpoint Blockade response. (A) Lethally irradiated C57Bl/6J were rescued with Tet2^wt^,Tet2^het^ or Tet2^KO^ bone marrow cells, followed by MC38 cell implantation and treatment with PD-1 ICB or isotype control antibody. (B) MC38 tumors are smaller in Tet2^het^ mice treated with ICB. Values from multiple unpaired t-tests comparing mean tumor volume between Tet2^het^ and Tet2^wt^, corrected for multiple hypothesis testing with Benjamini and Hochberg method. (C) MC38 growth curves for PD-1 treated mice with Tet2^wt^,Tet2^het^ or Tet2^KO^ bone marrow rescue from n=3 pooled experiments, with the fraction of complete responders achieving endpoint tumor volume <2 mm^3^ shown. p value from Fisher’s Exact Test versus Tet2^wt^. (D) CD45.1^+^ Tet2^wt^ versus CD45.1^+^ CD45.2^+^ Tet2^het^ bone marrow competition experiment in lethally irradiated recipient CD45.2^+^ B6 mice implanted with MC38 tumors and treated with PD-1 ICB. (E) The percent of CD45.1^+^ CD45.2^+^ Tet2^het^ (top) or CD45.1^+^ Tet2^wt^ donor-derived cells out of all cells for a given population in the bone marrow (BM), spleen, and MC38 tumors pre-treatment (light gray, n=4) and post-PD-1 ICB treatment (dark gray, n=5). Pre- and Post-treatment cell frequencies were compared by unpaired Student’s T Test. ns, *, ** represent p >0.05, p < 0.05, p < 0.01, respectively.

We then compared the effect of anti-PD-1 ICB in Tet2^wt^ and Tet2^het^ mice with established MC38 tumours. Tumour growth in anti-PD-1 treated Tet2^het^ mice was significantly attenuated compared to Tet2^wt^ mice (Figure 1B, Day 28 mean Tet2^het^ tumour volume 51.6% of Tet2^wt^, q value 0.004 from multiple unpaired t-tests). In three pooled experiments, we compared the likelihood of ICB-mediated control (endpoint tumor < 2 mm^3^) of newly implanted MC38 tumors in mice with Tet2^wt^, Tet2^het^, or Tet2-homozygous null (Tet2^KO^)-hematopoiesis. Tet2^het^ mice showed a trend towards greater tumor control with PD-1 blockade (Tet2^het^ vs Tet2^wt^ OR 3.1, p=0.129 Fisher’s exact test), with significantly greater odds of tumor control in immunotherapy-treated Tet2^KO^ compared to Tet2^wt^ mice (OR 6.8, p = 0.02, Figure 1C). Compared to Tet2^wt^ mice, endpoint anti-PD-1 treated tumors were smaller in Tet2^het^ (p = 0.020, Mann-Whitney Test) and Tet2^KO^ mice (p = 0.013), though tumor size did not differ between Tet2^het^ and Tet2^KO^ mice (Supplementary Figure 1D, p= 0.436). Tumor control rate and endpoint tumor volume did not significantly differ between isotype-control treated Tet2^wt^, Tet2^het^ and Tet2^KO^ mice (Supplementary Figure 1E and 1F). These results indicate that Tet2 inactivation in leukocytes potentiates anti-PD-1 ICB in mice, which supports the clinical association between CH and improved survival in colorectal cancer patients treated with ICB^10^ or conventional chemotherapy^11^, and is in keeping with enrichment of ICB-response genesets in colorectal cancer patients with TET2-CH^7^. There was an interesting trend towards a dose-responsive effect whereby complete Tet2^KO^ could more effectively curb tumor growth than heterozygous *Tet2* mutation. Since patients with TET2-CH most commonly have a single mutated *TET2* allele, we focused on Tet2^het^ and Tet2^wt^-rescued mice in subsequent experiments.

To understand how *Tet2* mutation alters hematopoietic dynamics and TIL frequency in response to anti-PD1 ICB we performed a competition experiment, where wild type CD45.2^+^ mice were rescued from lethal irradiation with a 50:50 mix of Tet2^het^ CD45.1^+^CD45.2^+^ and Tet2^wt^ CD45.1^+^ bone marrow. This allowed us to distinguish residual recipient mouse hematopoietic cells (CD45.2^+^) from donor-engrafted Tet2^het^ (CD45.1^+^CD45.2^+^-double positive) or Tet2^wt^ (CD45.1^+^) cells in recipient bone marrow, spleen, and tumours, before and after immunotherapy (Figure 1D, Supplementary Figure 1G-J). Tet2^het^ cells exhibited a significant competitive advantage and composed the majority of stem and progenitor cells in the bone marrow; this further increased over the course of anti-PD-1 treatment (Supplementary Figure 1G-1I). The proportion of Tet2^het^ dendritic cells, macrophages, and neutrophils was correspondingly high in tumours and spleen, and increased further during immunotherapy (Figure 1E, Supplementary Figure 1G and 1H). This is in keeping with the known repopulating advantage and myeloid bias of *Tet2*-mutant HSCs^13^. While nearly all common lymphoid progenitors were Tet2^het^, there was a lower fraction of Tet2^het^ lymphoid compared to myeloid cells in tumors and spleen (Figure 1E), reflecting myeloid bias of Tet2^het^ progenitors and the presence of residual recipient lymphocytes. The proportion of Tet2^het^ lymphoid TILs did not significantly increase with treatment (Figure 1E and Supplementary Figure 1H-1J). Therefore, the enhanced ICB response observed in mice with Tet2^het^ hematopoiesis correlates with significantly increased production of myeloid cell populations, without an appreciable effect on lymphopoiesis. The frequency of most Tet2^het^ TIL subpopulations was proportional to their frequency throughout the hematopoietic hierarchy. Besides dendritic cells, which became significantly enriched in post-treatment tumors and spleens but not bone-marrow, the results did not support a competitive advantage or enhanced tumor-infiltrating potential for mutant cells. Therefore, we reasoned that Tet2 inactivation may potentiate ICB by re-shaping TIL response to PD-1 blockade.

### *Tet2*-mutation promotes anti-tumor T cell states upon PD-1 blockade

To understand how somatic Tet2-mutations influence TIL frequency and alter TIL state with ICB, we performed single cell RNA-sequencing (scRNA-seq) with targeted TCR variable, diversity, joining (VDJ) analysis. We profiled 38,994 CD45^+^ TILs from size-matched isotype control or anti-PD-1 treated MC38 tumors in lethally irradiated mice with complete Tet2^wt^ or Tet2^het^ hematopoietic rescue (Figure 2A, Supplementary Figure 2A and 2B). Data integration, normalization, and batch correction were performed prior to clustering, and the resulting clusters of TIL subtypes were annotated using prior knowledge and by comparison to cell type-specific signatures from published scRNA-seq analyses of murine TILs^14,15^. With isotype-control treatment, Tet2^het^ TILs were enriched for B cells, naïve CD4 and naïve CD8 T Cells, and myeloid-derived suppressor cells (MDSCs), with relatively fewer mast cells and cycling NK cells (Supplementary Figure 2C and 2D). With PD-1 blockade, we observed the following in Tet2^het^ lymphocytes (red) compared to wt controls (black): CD4^+^ T cells were more likely to exhibit a naïve or memory phenotype and were less likely to be regulatory CD4^+^ T cells (Figure 2B) following ICB. CD8^+^ T cells were more likely to have a naïve or memory phenotype, while CD8^+^ T cell exhaustion was only repressed in Tet2^het^ TILs (Figure 2B). Thus, PD-1 blockade skewed the phenotype of Tet2^het^ T cells towards anti-tumor states compared to Tet2^wt^ TILs.

**Figure 2.**
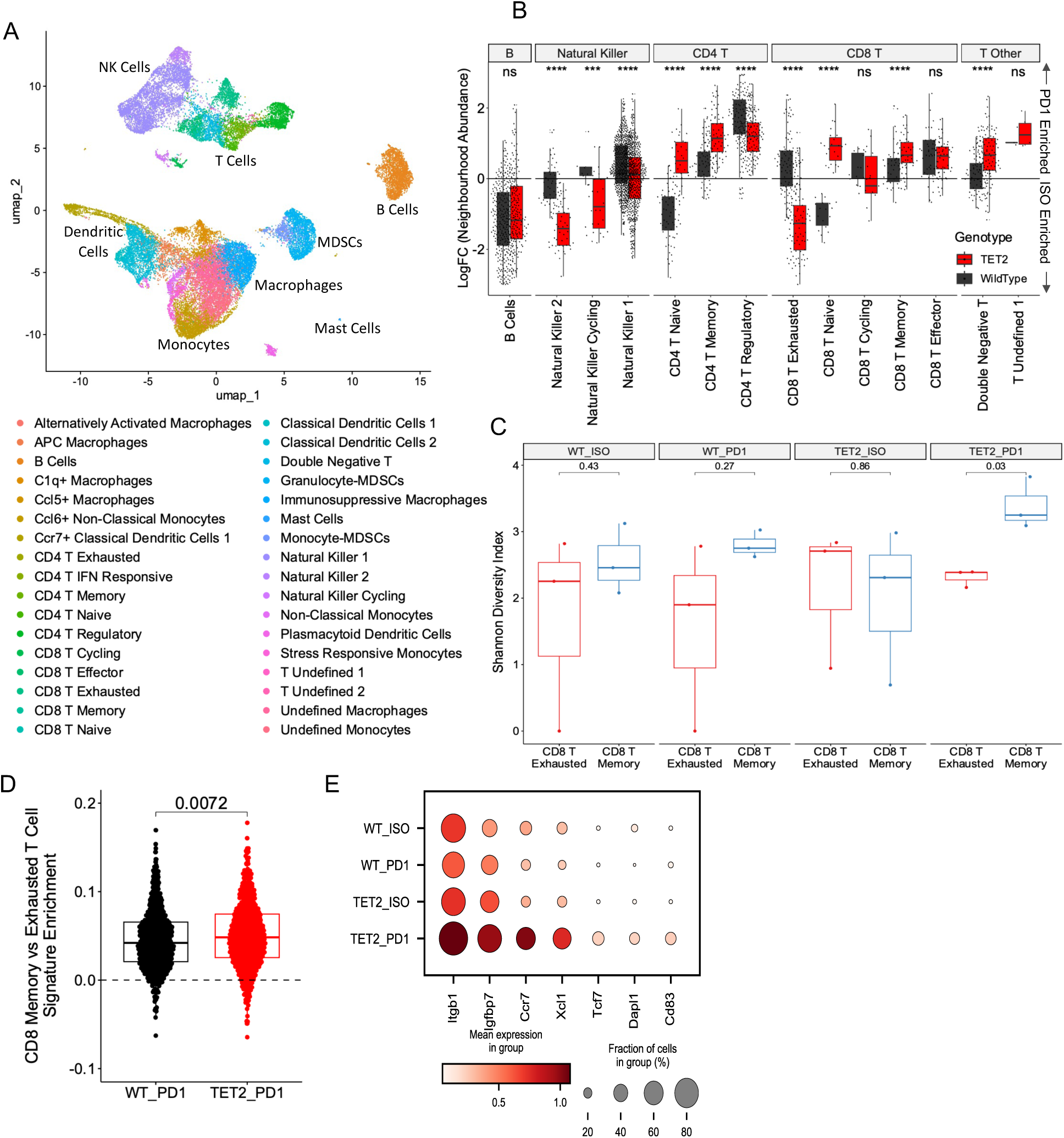
Tet2^het^ T cells are biased towards memory over exhausted or regulatory states with PD-1 ICB. (A) UMAP projection of annotated clusters identified in scRNA-seq from n=38,994 CD45^+^ TILs from MC38 tumors in mice with Tet2^wt^ or Tet2^het^ hematopoiesis following isotype control (ISO, n=3 Tet2^wt^, n=3 Tet2^het^) or anti-PD-1 treatment (PD1, n=5 Tet2^wt^, n=5 Tet2^het^). (B) Lymphoid TIL subtype abundance from Milo neighborhood analysis in PD1 relative to ISO-control treated tumors by mouse hematopoietic genotype (Tet2^wt^ = black, Tet2^het^ = red) displayed as log fold change from ISO to PD1 treatment. * = p<0.05, ** = p<0.01, *** = p<0.001, **** = p<0.0001, ns = not significant. (C) T cell receptor VDJ Shannon Diversity Index by mouse genotype and treatment from CD8 exhausted and CD8 memory T cells identified in scRNA-seq from n = 3 Tet2^wt^ ISO, Tet2^het^ ISO, Tet2^wt^ PD1, and Tet2^het^ PD1 mice. P values are shown from two-tailed, unpaired student’s T test. (D) Enrichment of the CD8 Memory T cell vs Exhausted T cell gene signature from GSE9650 compared with AUCell between Tet2^wt^ and Tet2^het^ anti-PD-1 treated CD8 T cells. p value from unmatched Mann-Whitney U Test. (E) For genes significantly upregulated in anti-PD-1 treated Tet2^het^ (TET2) vs Tet2^wt^ (WT) CD8 Exhausted cells, dot plot of mean gene expression and fraction of cells expressing each gene in CD8 Exhausted WT and TET2 cells across ISO and PD1 treatment.

Tet2-loss of function promotes memory CD8 gene expression programs, which may bias antigen-stimulated cells towards this state at the expense of exhaustion^6,16^. To evaluate this, we quantified clonal diversity of T cells through VDJ recombination analysis. We found that T cell clonal diversity was greater in Tet2^het^ anti-PD-1 treated mice compared to isotype control or Tet2^wt^ PD-1 treated mice (Supplementary Figure 2E). Interestingly, higher TCR diversity has been linked to ICB response in melanoma patients^17^. Importantly, memory CD8 T cell diversity was greater than exhausted CD8 T cell diversity in immunotherapy-treated Tet2^het^ mice but not other conditions (Figure 2C). Greater diversity in the memory CD8 pool reflects mutant T cell priming for memory formation at the expense of exhaustion upon the persistent TCR signaling that occurs during ICB. Consistent with this possibility, CD8 T cells in Tet2^het^ mice showed enrichment of genes associated with a memory CD8 state relative to exhausted state (Figure 2D). In keeping with the ability for Tet2-loss to promote the memory CD8^+^ T cell phenotype, genes related to memory CD8 programs persisted throughout T cell activation states and remained upregulated in Tet2^het^ versus Tet2^wt^ cells within the exhausted CD8 cluster (Figure 2E).

*Tet2* transcription is activated following TCR engagement and serves to suppress T cell effector and memory functions in mouse models and adoptive cell therapy^5,6,16^. Our animal model data suggest that Tet2-mutant CD8 T cells are biased towards memory and effector states and suppress regulatory and exhausted states with immunotherapy. TET2-CH is a risk factor for T cell lymphoma, and prior studies have detected *TET2* CH mutations in T cells, albeit at lower variant allele frequency (VAF) than myeloid populations and B cells^18,19^. There are reports of *TET2*-mutant T cells undergoing dramatic expansion in CAR-T^6^ or ICB^20^-treated patients who respond to immunotherapy. Therefore, harnessing the positive aspects of TET2-inactivation while avoiding uncontrolled lymphocyte proliferation or comorbid inflammatory sequalae may have therapeutic potential.

### Tet2 inactivation shifts myeloid cells from immunosuppressive to costimulatory states with PD-1 blockade

We further investigated the impact of *Tet2* mutations on the response of myeloid TILs to ICB and myeloid-T cell interactions within tumors. Notably, PD-1 treated Tet2^het^ tumors contained significantly more cells within classical dendritic cell neighborhoods (Figure 3A). Both granulocyte- and monocyte-MDSCs were significantly more abundant in Tet2^het^ tumors (Figure 3A). However, differential gene expression analysis revealed upregulation of T cell-activating molecules *Cxcl9* and *Ccl5*, and downregulation of immunosuppressive *Ccl2*, in Tet2^het^ MDSCs (Figure 3B). Immunosuppressive macrophage fate was also induced by PD-1 blockade in Tet2^wt^ controls but this was attenuated in Tet2^het^ tumors (Figure 3A). Together with the MDSC findings, this is consistent with prior reports of *Tet2*-mutant myeloid TILs being less immunosuppressive than Tet2^wt^ myeloid TILs^4^. While the relative abundance of monocyte subtypes varied considerably, the gene expression programs enriched in *Tet2*-mutant monocytes were related to T cell activation through co-stimulation (Figure 3C), including genesets whose enrichment is associated with immunotherapy response in murine and human tumours^21^. DNA methylation analysis in F4/80^+^ monocyte/macrophage TILs from isotype and anti-PD-1 treated tumors revealed hypermethylation of probes within genes related to immune activation and myeloid cell differentiation (Figure 3D, Supplementary Figure 3A). Promoters of genes defining the immunosuppressive macrophage phenotype were hypermethylated in Tet2^het^ cells (Supplementary Figure 3B). Therefore, DNA hypermethylation due to Tet2 loss of function is linked to differences in cell state induced by ICB. This suggests that *Tet2* mutant leukocytes have a unique epigenetic state that controls the response to immunotherapy.

**Figure 3.**
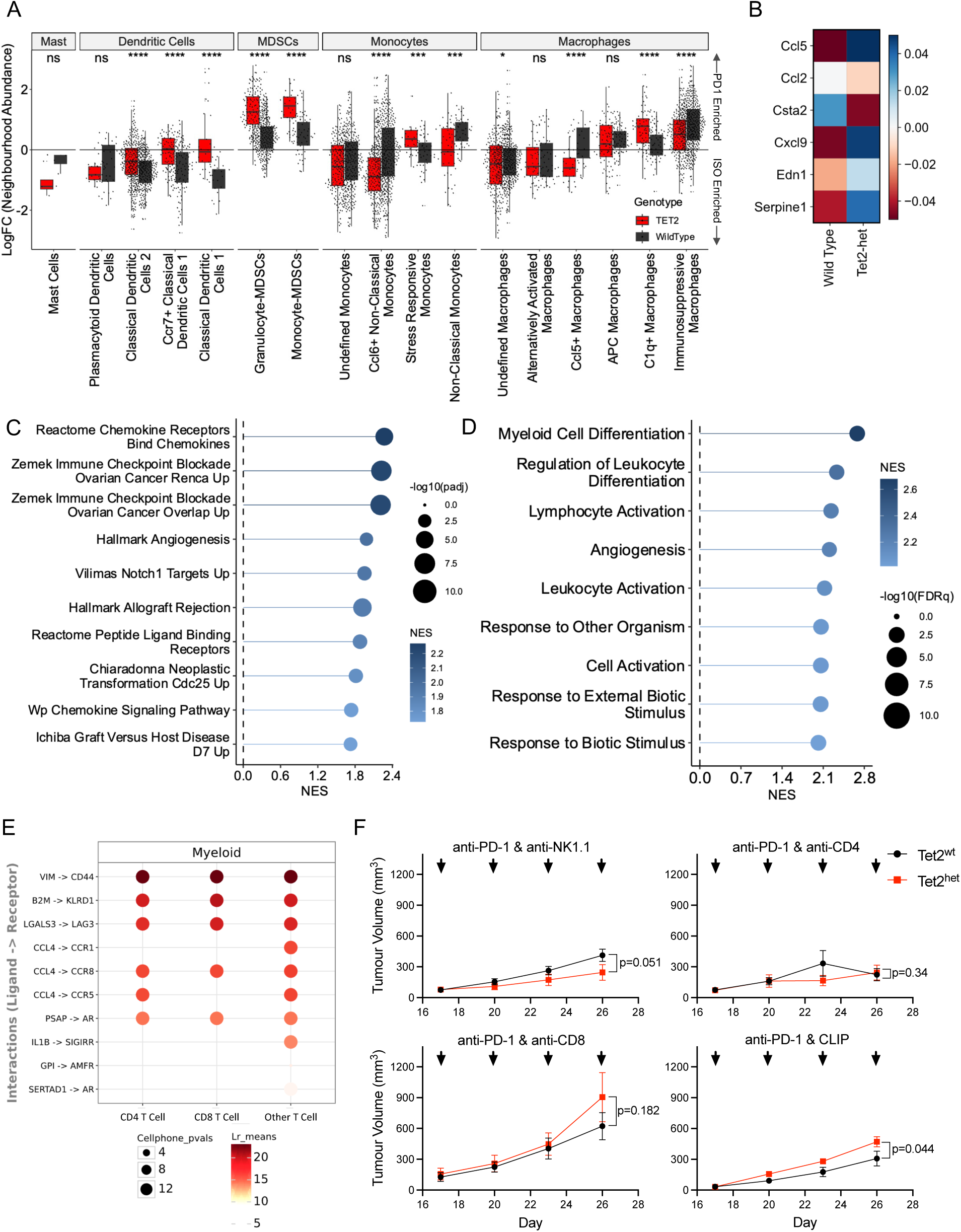
Tet2^het^ mutant phagocytes, CD4, and CD8 T cells are required to promote PD-1 response (A) Myeloid TIL subtype abundance from Milo neighborhood analysis in PD1 relative to ISO control treated tumors by mouse hematopoietic genotype (Tet2^wt^ = black, Tet2^het^ = red) displayed as log fold change from ISO to PD1 treatment. * = p<0.05, ** = p<0.01, *** = p<0.001, **** = p<0.0001, ns = not significant. (B) Genes significantly differentially expressed (FDR<0.05 per DESeq2) in Tet2^het^ versus Tet2^wt^ anti-PD-1 treated MDSCs by scRNA-seq. (C) Gene sets significantly enriched in Tet2^het^ compared to Tet2^wt^ PD-1 treated monocytes by GSEA. (NES = normalized enrichment score). (D) Gene Ontology (GO) enrichment terms from hypermethylated probes in distal cis-regulatory domains hypermethylated in Tet2^het^ versus Tet2^wt^ F4/80^+^ cells from isotype control and anti-PD-1 treated MC38 tumors. Significantly enriched terms with probes detected in at least 10 genes are shown. (E) LIANA consensus cell-cell communication analysis between myeloid (sender) and CD4, CD8, or other T cell populations (receiver), showing interactions significantly enriched in Tet2^het^ PD-1 treated compared to Tet2^wt^ PD-1 treated TILs with p values from CellphoneDB represented as circle diameter. (F) Mean MC38 tumor volume in mice rescued with Tet2^wt^ (black) or Tet2^het^ (red) hematopoiesis treated with PD-1 ICB plus NK-cell (n=10), CD4-(n=7), or CD8-depleting antibodies (n=7), or clodronate liposomes (CLIP, n=6). Mice were matched based on day 17 tumor volume. Endpoint p values from one-tailed Student’s T test.

It is unclear how much the observed increases in T cell activation/memory formation, and decreases in exhaustion, can be attributed to T cell-intrinsic effects of mutations versus altered co-stimulation from cells in the microenvironment. Ligand-receptor analysis framework (LIANA)^22^ based cell-cell communication analysis of scRNA-seq data showed increased T cell – antigen presenting cell interactions and enhanced T cell stimulation from myeloid cells through CD44, CCR5, and the TCR in Tet2-mutant tumors (Figure 3E). Therefore, while differential gene expression in mutant monocytes and T cells revealed a degree of cell-autonomous priming, it is likely that a combination of altered interactions and costimulatory signals from myeloid cells in the tumor microenvironment, plus cell-intrinsic rewiring from Tet2-loss, promotes memory T cell fates and limits exhaustion or regulatory fate upon PD-1 blockade. Thus, the improved ICB response with Tet2 inactivation is likely due to simultaneous cell autonomous and non-cell autonomous effects.

### Enhanced ICB response with Tet2 inactivation requires T cells and phagocytes

To identify the TIL subtypes necessary to improve ICB response with Tet2-mutation, we performed targeted cell depletion in anti-PD-1 treated mice using blocking antibodies or clodronate liposomes. Tet2^het^ mice still showed enhanced anti-PD-1 response following depletion of natural killer (NK) cells (n=10 per group, endpoint Tet2^het^ tumor volume 60% of Tet2^wt^, p = 0.051 by unpaired T-test, Figure 3F, Supplementary Figure 3C). Depletion of CD4^+^ T cells or CD8^+^ T cells with CD4- or CD8-targeted antibodies, respectively, eliminated the Tet2^het^ therapeutic effect and led to comparable tumor growth kinetics between Tet2^wt^ and Tet2^het^ anti-PD-1 treated mice (n = 7 per group, CD4 depletion: Tet2^het^ tumor volume 107% of Tet2^wt^, p = 0.34; CD8 depletion: Tet2^het^ tumor volume 145% of Tet2^wt^, p = 0.34, Figure 3F, Supplementary Figure 3D and 3E). Depletion of phagocytes, including macrophages and dendritic cells with clodronate liposomes, not only eliminated the Tet2^het^ therapeutic effect, but led to significantly larger endpoint tumors in anti-PD-1 treated Tet2^het^ versus Tet2^wt^ mice (n=6 per group, Tet2^het^ tumor volume 153% of Tet2^wt^, Figure 3F, Supplementary Figure 3F and 3G). These findings indicate that hematopoietic loss of Tet2 function promotes ICB response through T cells and phagocytes, but not NK cells.

### TET2-CH is associated with immune infiltration and enhanced clinical benefit from immune checkpoint blockade in melanoma

To determine whether our murine modeling was clinically relevant, we focused on human melanoma, as this is an archetypal ICB-responsive cancer in which to explore the consequences of TET2-CH on tumor biology and response to immunotherapy. To assess TET2-CH-induced changes in the melanoma tumor microenvironment, we paired bulk RNA-sequencing analysis with TET2-CH calls in the TCGA SKCM cohort, identifying patients with CH by detecting somatic mutations in peripheral blood exome data (Methods). Compared to patients without detectable CH mutations, tumors from TET2-CH patients showed enhanced immune infiltration by CIBERSORT_x_ (Supplementary Figure 4A). Specifically, TET2-CH tumors were enriched for T cells, dendritic cells, and B cells (Figure 4A-C). GSEA comparing tumors from TET2-CH^+^ patients to patients without CH revealed strong enrichment for innate and adaptive immune activity, including greater T cell receptor (TCR) signaling in patients with TET2-CH (Figure 4D and Supplementary Figure 4B). Bulk melanoma samples from TET2-CH^+^ patients showed upregulation of the genes defining a T cell centric - dendritic cell enriched immune microenvironment that is associated with improved survival in melanoma, and solid tumors in general (Figure 4E)^23^. Chronic antigen stimulation and TCR signaling in infection or cancer is tolerated in memory T cells, but can also lead to T cell exhaustion. Melanomas from patients with TET2-CH showed a greater memory CD8 T cell versus exhausted CD8 T cell signature (Figure 4F). These results suggest that TET2-CH is associated with greater immune infiltrate, adaptive immune activation and less T cell exhaustion, in untreated melanoma.

**Figure 4.**
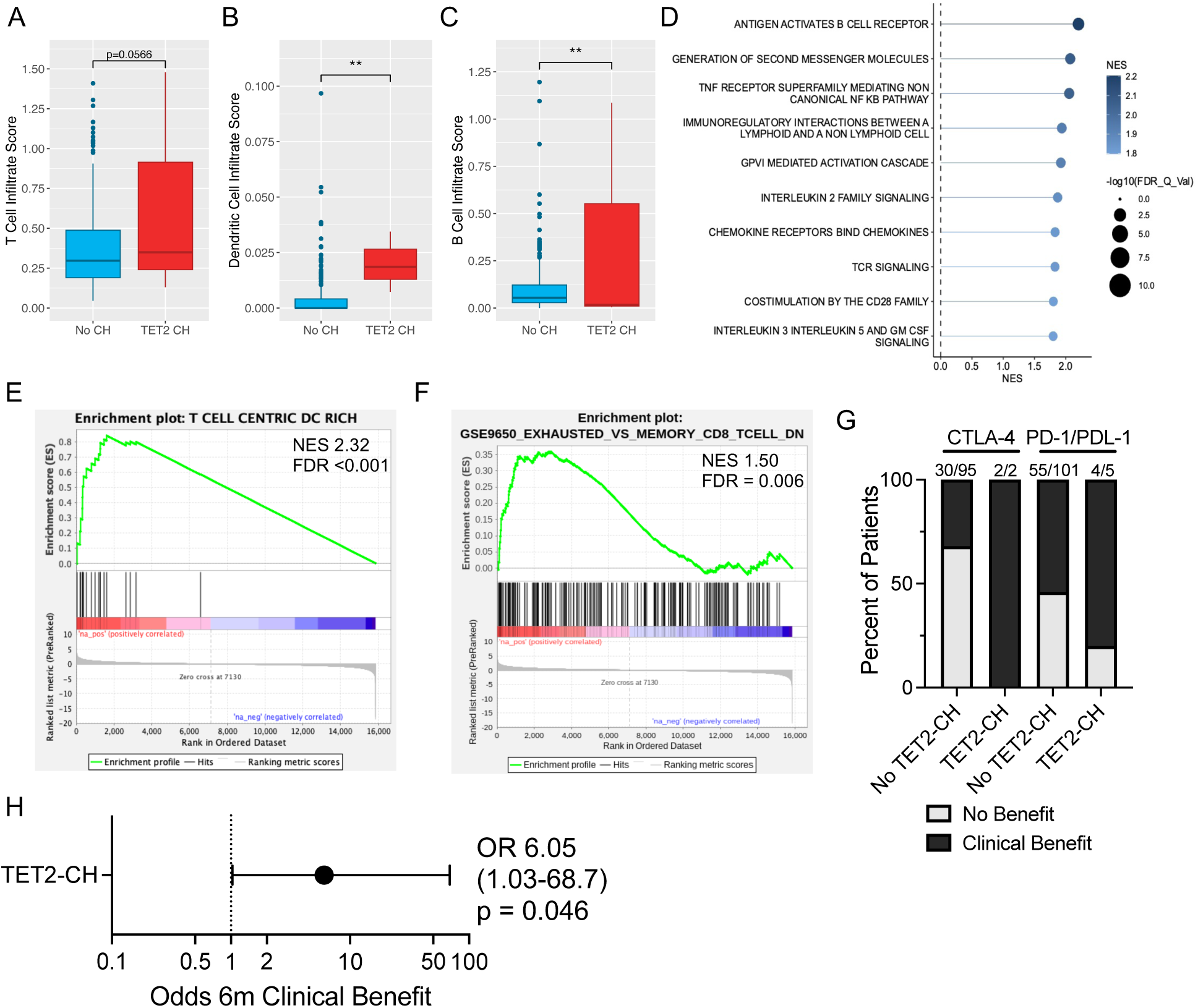
TET2-CH is associated with clinical benefit from ICB in melanoma patients. (A-C) CIBERSORT_x_ T cell (A), Dendritic Cell (B), and B Cell (C) infiltrate score in TCGA-SKCM melanoma bulk RNA-sequencing from patients with TET2-CH and without CH compared by Mann-Whitney U test. (D) Normalized enrichment score and -log10FDRq value from GSEA analysis of bulk RNA-sequencing comparing Reactome geneset enrichment in TET2-CH positive Melanoma tumors to CH negative Melanoma tumours. (E) GSEA enrichment plot with normalized enrichment score (NES) and false discovery rate (FDR) for the T Cell Centric DC Rich tumor immune microenvironment signature in melanoma tumors from patients with *TET2*-CH versus without CH. (F) GSEA enrichment plot with normalized enrichment score (NES) and false discovery rate (FDR) for genes downregulated in exhausted CD8 T cells versus memory CD8 T cells (GSE9650) in melanoma tumors from patients with *TET2*-CH versus without CH. (G) Percent of metastatic melanoma patients deriving clinical benefit (gray) or no clinical benefit (black) from CTLA-4 or PD-1/PDL-1 targeted ICB at 6 months. Number of responders and number of patients assessed is shown above each column. (H) Odds ratio of 6-month clinical benefit in melanoma patients exposed to TET2-CH versus no TET2-CH (G) with 95% confidence interval, shown by primary cancer type, using Firth’s penalized logistic regression with age, sex, study, and immune Checkpoint as covariates. *, ** denotes p<0.05, p<0.01, respectively.

To test for an association between the presence of *TET2*-mutant CH and immunotherapy response, we re-analyzed sequencing data from 203 ICB-treated melanoma patients across 4 studies to identify those with TET2-CH mutations detectable in exome sequencing data (Supplementary Figure 4C, Methods, Supplementary Table 1)^24–27^. 97 patients were treated with anti-Cytotoxic T lymphocyte associated protein 4 (CTLA-4) ICB and 106 patients were treated with ICB targeting PD-1 or its ligand Programmed death-ligand 1 (PD-L1). Patients with CH mutations were significantly older than those without (Supplementary Figure 4D). We assessed outcomes following ICB at 6 months, with radiographic response, clinical response, or stable disease defined as clinical benefit, with lack of clinical benefit defined by clinical progression, radiographic progression, or death (Figure 4G). When compared to patients without TET2-CH in multivariate analysis, the three percent of metastatic melanoma patients with TET2-CH were 6-fold more likely than non-TET2-CH cases to derive clinical benefit from ICB when adjusting for covariates patient age, sex, study, and immune checkpoint (OR 6.05 [95%CI], p = 0.046 multivariate logistic regression, Figure 4H). As reported in a previous study of melanoma patients, CH with any driver mutation was not associated with clinical benefit from ICB (Supplementary Figure 4E).

In summary, we demonstrate that Tet2 inactivation reshapes the tumor-immune microenvironment by priming costimulatory and memory programs in myeloid and T cells, respectively, and this is associated with greater clinical benefit in melanoma patients. We show how somatic mutations originating in HSCs can alter solid tumor biology with clinical implications. Future studies are required to determine the underlying mechanisms and explore how to translate these findings to improve cancer patient outcomes.

## ACKNOWLEDGEMENTS

We would like to thank the Sickkids/UHN and DKFZ Flow Cytometry Facilities, the UHN Animal Resource Centre, all patients and their families. Figures 1A, 1D, and Supplementary Figure 2A were created with BioRender.

## METHODS

### Mouse bone marrow transplant model

Female C57Bl6/J mice (JaX Strain 000664, RRID: IMSR_JAX:000664) or CD45.1 C57Bl6/J (JaX Strain 0002014, RRID: IMSR_JAX:0002014) between 6-10 weeks of age were lethally irradiated with two doses of 6 Gy from a Cesium source administered 4 hours apart. Mice were maintained on Enrofloxacin for 14 days beginning on the day of irradiation. One day post-irradiation mice were rescued by tail-vein injection of 1-1.5 million bone marrow cells from female donor mice, aged matched between 8-16 weeks of age, that lacked (Tet2^wt^, JaX Strain 000664, RRID: IMSR_JAX:000664 or JaX Strain 0002014, RRID: IMSR_JAX:0002014), were heterozygous (Tet2^het^, Jax Strain 023359; RRID: IMSR_JAX:023359 alone or crossed to JaX Strain 0002014, RRID: IMSR_JAX:0002014 CD45.1 mouse) or homozygous (Tet2^KO^) for a deletion in Tet2 exons 8-11, including the catalytic domain, which leads to a loss of detectable gene product in homozygotes (Jax Strain 023359; RRID: IMSR_JAX:023359). Equal cell numbers were used for each genotype of donor mouse in an experiment.

### Syngeneic tumour implantation and immunotherapy treatment

9 weeks after bone marrow transplant, 250,000 MC-38 cells (Kerafast, Shirley, USA, RRID: CVCL_B288) were subcutaneously implanted into the right flank of rescued mice. MC-38 cells were below passage 8 and tested negative for mycoplasma or other contaminants. At the time of tumor implantation, mice were randomized to treatment with 300 μg of anti-PD-1 blocking antibody (BioXcell clone RMP1-14, RRID: AB_10949053) or rat IgG2a isotype control (BioXcell clone 2AE, RRID: AB_1107769) delivered by intra-peritoneal injection once every 3 days, beginning on day 10 post-implantation for a total of 4 doses. Tumor volume was measured using the formula volume = ½(length x width^2^). For antibody-based cell depletion experiments, rescued mice were implanted with 250,000 MC-38 cells in the right flank and at Day 17 tumor volume was measured using the formula volume = ½(length x width^2^). Mice rescued with Tet2^wt^ and Tet2^het^ bone marrow were then matched by tumor volume and assigned to intra-peritoneal treatment with 300 μg of anti-PD-1 blocking antibody (BioXcell clone RMP1-14, RRID: AB_10949053) with or without cell depleting antibody targeting CD4 (100 μg BioXcell clone GK1.1, RRID: AB_1107636), CD8 (100 μg BioXcell clone 2.43, RRID_1125541), or NK1.1 (25 μg BioXcell clone PK136, 1107737) delivered every 3 days for 4 doses. Liposomal clodronate depletion of macrophages by intraperitoneal injection of 1.5 mg then 1 mg doses were administered in parallel to 300 μg of anti-PD-1 blocking antibody (BioXcell clone RMP1-14, RRID: AB_10949053) every 3 days for four total doses to mice rescued with Tet2^wt^ or Tet2^het^ bone marrow, having size-matched tumors from implantation of 125,000 MC-38 cells in the right flank. Investigators were not blinded to the experimental groups.

At pre-specified endpoints tumors were dissected and dissociated with 1X Collagenase and Hyaluronidase in DMEM (Stemcell Technologies, Vancouver, Canada) with 0.1 mg/mL DNAse (Roche, Basel, Switzerland) for 1 hour at 37 C. Fragments were then washed in DMEM with 2% fetal calf serum, filtered through 70 μm and 40 μm filters and subjected to 5 minute ammonium chloride lysis at room temperature prior to use in downstream applications.

For BM competitive transplantation experiments, mice were euthanized and spleen, injected femur, and tumor were collected. The condyle of the femur was removed, and cells harvested by centrifugation. Spleen was gently mashed using a plunger. Tumors were dissociated as above. Cell collection was performed in PBS FBS 2% and filtered through a 35-µm nylon strainer prior evaluation by flow cytometry. For analysis of mature cell populations, cells were stained with the following Abs: CD45.1-BV421 (RRID_ AB_10896425), CD45.2-APC (RRID_ AB_469400), B220-APC/Cy7 (RRID_ AB_313006), CD3-PE (RRID_ AB_312662), CD11b-APC/Cy7 (RRID_ AB_312788), CD11c-PeCy7 (RRID_ AB_493569), F4/80-PE (RRID_AB_2687527) and Ly6G-FITC (RRID_ AB_2562352). For analysis of HSPCs, cells were stained with the following Abs: CD45.1-APC/Cy7 (RRID_ 2629806), CD45.2-Alexa Fluor 700 (RRID_ AB_493731), Lineage-FITC (RRID_ AB_11150779), CD48-APC (RRID_ AB_571997), CD117-PE (RRID AB_394806), CD127-PeCy5 (RRID_ AB_1937262), CD135-PeCy7 (RRID_AB_2737639), CD150-BV711 (RRID_ AB_2629660) and Sca-1-BV421 (RRID_AB_ 2563064). Viability was assessed using Propidium Iodide. Analysis was performed using the BD FACSSymphony and FlowJo.

### Mouse tumor-infiltrating leukocyte single cell RNA-sequencing

9 weeks after rescue with Tet2^wt^ or Tet2^het^ bone marrow cells as described above, C57Bl6/J mice (JaX Strain 000664, RRID: IMSR_JAX:000664) were subcutaneously implanted with 250,000 MC-38 cells (Kerafast, Shirley, USA, RRID: CVCL_B288) in the right flank. 17 days post-tumor implantation Tet2^wt^ or Tet2^het^ rescued mice were matched by tumor-size and assigned to receive 300 μg of anti-PD-1 blocking antibody (BioXcell clone RMP1-14, RRID: AB_10949053) or rat IgG2a isotype control (BioXcell clone 2AE, RRID: AB_1107769) every 2 days for 3 doses. Mice were euthanized 48 hours after the final dose and tumors processed as described above. Dead cells were removed by negative selection using the EasySep Dead Cell Depletion Kit (Stemcell Technologies, Vancouver, Canada). Cells were stained for CD45 (BD Biosciences clone 30-F11, RRID: AB_394609) and with Sytox Blue (ThermoFisher, Waltham, USA) to identify remaining dead cells. Live, CD45^+^ tumor-infiltrating leukocytes were isolated using Fluorescence-Activated Cell Sorting with a Sony MA900 device (Sony Biotechnology, San Jose, USA). Single cells were processed using Chromium 10X 5’ Single Cell and Chromium Single Cell Mouse TCR Amplification kits at the Princess Margaret Cancer Centre Genomics Facility according to the manufacturer’s instructions (10X Genomics, Pleasanton, USA).

Sequencing data for 38,994 total cells from isotype control (n=3 Tet2^wt^ and n=3 Tet2^het^) and anti-PD-1 treated mice (n=5 Tet2^wt^ and n=5 Tet2^het^) were analyzed using Seurat v4.3^28^ in the R programming environment^29^. The SCTransform method was used with “glmGamPoi” for data normalization and correction of technical variation, with principal component analysis applied for dimensionality reduction. Data was projected in to 2-dimensional space using the UMAP algorithm, and cells were grouped in to 20 clusters using Seurat’s FindNeighbors and FindClusters functions. Clusters were annotated based on prior knowledge, published gene expression signatures from MC38 TILs^14^, and similarities to human LM22 leukocyte gene expression programs. We projected our dataset onto the MetaTiME platform which provides pretrained meta-components from 1.7 million single cells across 79 tumor datasets to automatically annotate fine-grained cell states and plot signature continuum across our dataset.

To identify differentially enriched cell neighborhoods between PD-1 treated relative to ISO-treated tumors by mouse hematopoietic genotype, we performed Milo differential abundance analysis^30^. Cells were assigned to partially overlapping neighborhoods using a k-nearest neighbor graph using a neighborhood size of n=30. We performed multiple hypothesis correction following a weighted FDR procedure that accounts for spatial overlapping of the neighborhoods, as recommended by the Milo framework. Applying a spatial FDR cutoff of 0.1, we identified cell neighborhoods that were differentially enriched between our experimental conditions. AUcell was used to score gene expression signatures within neighborhoods between different conditions.

To quantify clonal diversity of CD8 gene expression programs, we performed VDJ recombination analysis using the R entropy package to calculate the Shannon Diversity Index of Tet2^wt^ and Tet2^het^ CD8 exhausted and CD8 memory T cells from isotype control or anti-PD-1 treated MC38 tumors.

We applied the consensus-based LIANA^22^ framework to estimate ligand-ligand interactions for sender cell populations, comprising cells in myeloid neighborhoods (macrophages, monocytes, dendritic cells, and MDSCs) and receiver T cell populations: CD4, CD8, or other T cell neighborhoods, to identify interactions significantly enriched in Tet2^het^ PD1-treated compared to TET2^wt^ PD1-treated TILs.

### Mouse tumor-infiltrating monocyte/macrophage DNA methylation analysis

Single cell suspensions were generated from isotype control or anti-PD-1-treated MC-38 tumors as described above. Viable, propidium iodide-negative (ThermoFisher, Waltham, USA), CD45^+^ F4/80^+^, CD3^-^ cells were isolated using fluorescence activated cell sorting with BD FACSAria II (Franklin Lakes, USA). DNA was isolated using the QiaAMP Micro Kit (Qiagen, Germantown, USA) and profiled using the Infinium Mouse Methylation BeadChip (Illumina, San Diego, USA). IDAT files were imported into RnBeads and subjected to quality control, background subtraction, and data normalization as previously described and using probe annotations described in Schönung et al. 2022^31^. Differentially methylated probes (DMPs) were identified between Tet2^het^ and Tet2^wt^ cells from Isotype control or anti-PD-1 treated tumors in a pairwise fashion using RnBeads “rnb.execute.computeDiffMeth” function, with a mean methylation difference of 10% and false-discovery-rate adjusted p value of <0.1. The overlap between genomic region class and DMPs was performed using annotations from Schönung *et al*. 2023 as previously described^31^. Gene Ontology (GO) terms associated with the genes possessing hypermethylated distal cis-regulatory domains (DCRD) in tumour-infiltrating Tet2^het^ monocyte/macrophages were identified using pathway analysis with g:Profiler^32^.

### Immune Checkpoint Blockade Response Analysis from Public Datasets

Fastq files for tumors and blood-derived ‘normal’ DNA were obtained from the following studies of immune checkpoint blockade response: Roh et al. 2017 (SRP115658)^24^, Hugo et al. 2016 (SRP067938, SRP090294)^25^, Snyder et al. 2014 (SRP072934)^26^, and Riaz et al. 2017 (SRP095809)^27^. Data was aligned to GRCh38 with bwa v0.7.17 and collapsed with picard v2.21.4. Mutations were called using Mutect2 v4.1.2 and filtered using gatk v4.1.2 FilterMutectCalls. The mutation calls were processed with snpEff v5.0e. Coding region variants were filtered for frequencies of < 0.01 in gnomad and a minimum depth of 10. Variants were annotated as pathogenic based on criteria set out by Lindsley *et al*^33^. Finally for variants identified as pathogenic we screened for their presence in the matched tumor. We retained mutations where the ‘normal’ VAF was at least 2x higher than the VAF found in the tumor sequencing reads. Pathogenic variants passing the above filters at VAF ≥0.02 were considered CH variants, and patients were annotated as positive for CH accordingly.

To unify heterogeneous follow up data, 6 month clinical outcome was used as the endpoint. A patient was determined to have experienced clinical benefit from ICB if they remained alive and had a radiographic response (partial or complete) or stable disease for at least 6 months. A patient was deemed not to have benefited from ICB if they experienced tumor growth (typically 20% or more growth per RECIST1.1 or irRECIST) or died within 6 months. Patients without clinical outcome data were not included in the analysis. The odds of experiencing clinical benefit with/without exposure to any CH mutation or TET2-CH were calculated using Firth’s penalized multivariate logistic regression with patient age, sex, study, and immunotherapy treatment (PD-1 or CTLA-4 blockade) as covariates in the R programming environment.

### CHIP Detection in TCGA SKCM

Peripheral blood whole exome sequencing (WES) bam files aligned to the GRCh38.d1.vd1 reference genome from 468 TCGA participants with primary or metastatic melanoma were accessed via the Genomic Data Commons (GDC) portal.^34^ WES files were assessed for somatic variants using the Genome Analysis Toolkit (GATK)-Mutect2 algorithm. To overcome known issues with alignment and variant calling in *U2AF1*, samples were assessed for *U2AF1* variants using a custom pileup region script.^35^ Vcf outputs from GATK-Mutect2 were annotated using ANNOVAR for subsequent filtering.^36^

CHIP variant calling was performed following previously described best practices for large WES datasets.^35^ Briefly, somatic variants consistent with CHIP were called following a curated list of CHIP driver mutations in 67 genes. Variants were filtered to meet several sequencing quality metrics including coverage depth ≥20 reads, minimum alternative allele depth ≥5 reads, and F1R2/F2R1 ≥1 read. Variants with VAF below the 2% CHIP threshold were excluded, as well as variants with binomial test p >0.01, indicating a high likelihood of being a germline heterozygous variant. Indel variants occurring in homopolymer regions ≥3 base pairs and other variants recurring more frequently than known CHIP hotspots *DNMT3A* R882H/C and *JAK2* V617F were excluded as probable sequencing artefacts.

### TCGA SKCM Bulk RNA-seq Analysis

Aligned RNA-seq bam files from 437 primary and metastatic tumour samples with no known prior treatment exposures were accessed from the GDC portal. Files were de-aligned to fastq format using samtools, then reads were trimmed and filtered with fastp.^37^ Transcript pseudo-alignment and quantification was conducted using kallisto.^38^ Quality control checks were conducted with PCA plotting and MultiQC.^39^

Differential gene expression analysis was performed with DESeq2 to compare samples with *TET2*-mutant CHIP to those without CHIP.^40^ Analyses were controlled for sex and primary/metastatic tumour biopsy. Genes with p_adj_ <0.05 were differentially expressed. Preranked gene set enrichment analysis (GSEA) was performed to evaluate enrichment of Gene Ontology (GO) terms, HALLMARK pathways, and REACTOME pathways, as well as MSigDB exhausted vs memory CD8+ T cells and the top 10 differentially enriched pathways observed in the murine Tet2^het^ monocytes.^41^ Pathways demonstrating a false discovery rate (FDR) <0.1 were considered significantly enriched/supressed.

Immune cell subpopulations within the melanoma tumour microenvironment were deconvoluted via CIBERSORTx in absolute mode with the LM22 gene signature matrix.^42^ Differences in immune cell infiltrates between samples with *TET2*-mutant CH and no CH were evaluated using a generalized linear model adjusted for sex and primary/metastatic tumour biopsy. Only samples with high confidence (p <0.05) deconvolution were included in analyses.

## DATA AVAILABILITY STATEMENT

Single cell RNA-seq and DNA methylation array data will be made available through Gene Expression Omnibus (accession number pending).

