## Supplementary_Figures for "Hematopoietic Tet2 inactivation enhances the response to checkpoint blockade immunotherapy"

### Supplementary Figure 1

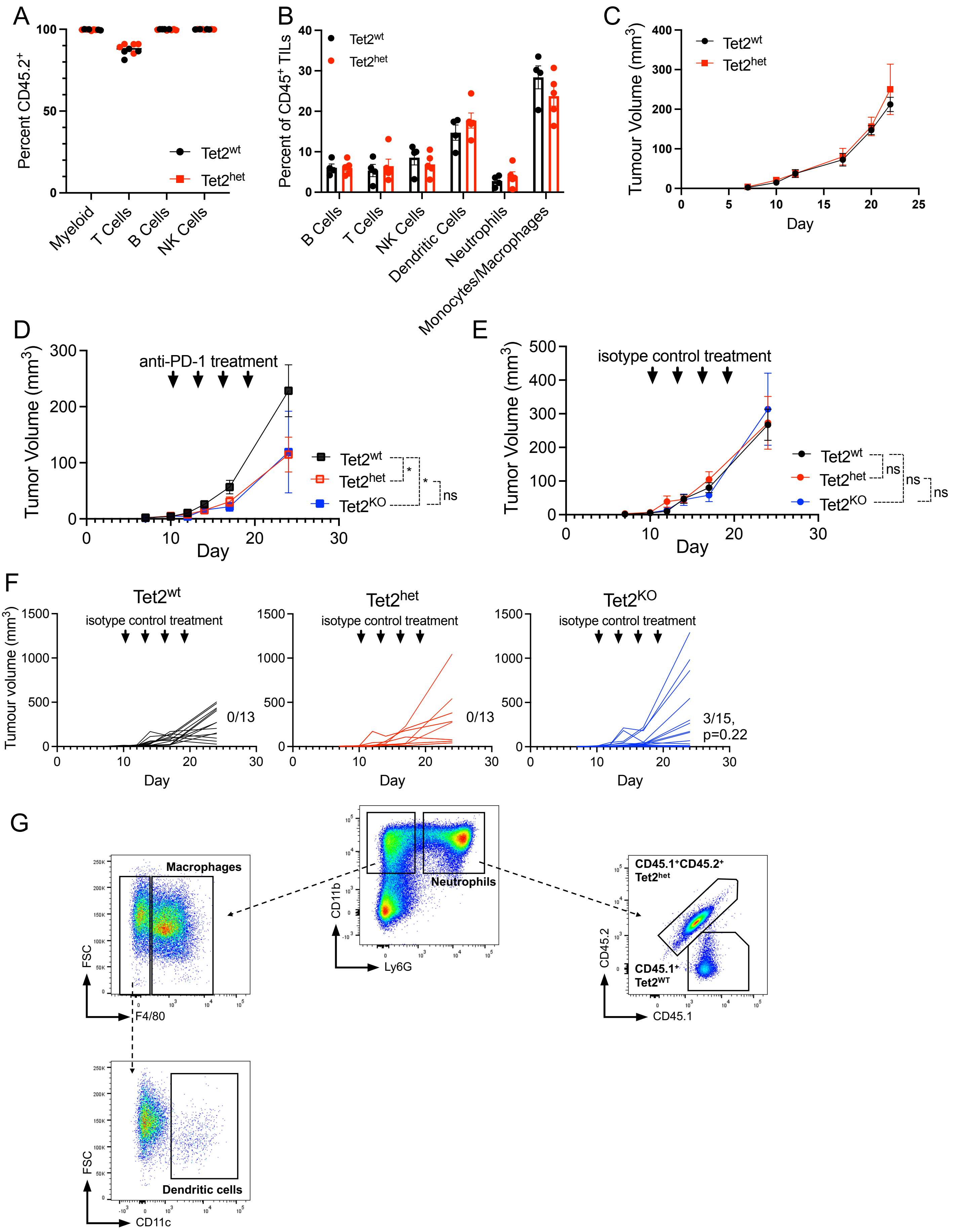

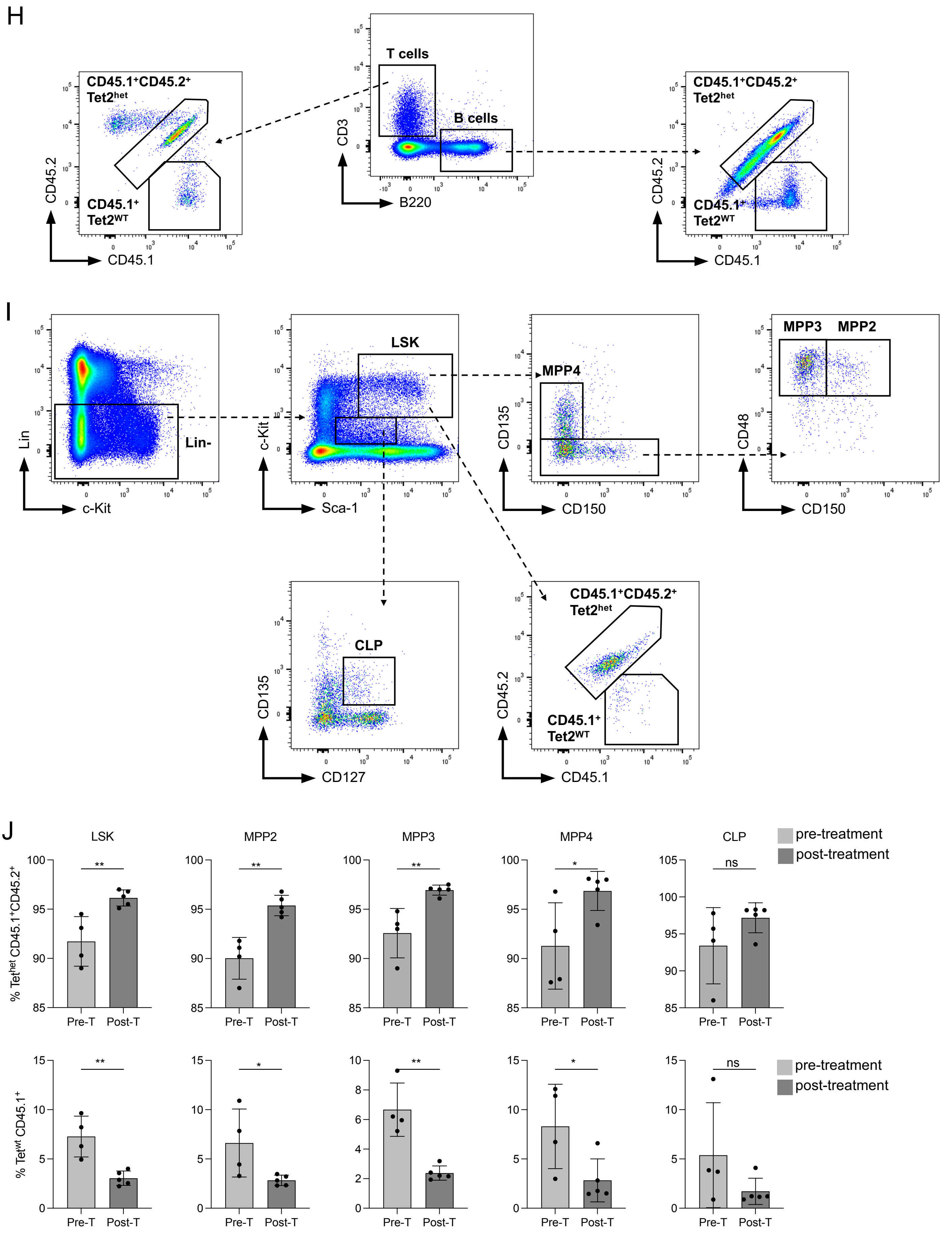
H
CD45.1+CD45.2+
Tet2het
CD45.1+
Tet2WT
CD45.2
CD45.1
T cells
B cells
CD3
B220
I
Lin-
Lin
c-Kit
LSK
Sca-1
MPP4
CD135
CD150
MPP3
MPP2
CD48
CLP
CD127
J
LSK
MPP2
MPP3
MPP4
CLP
pre-treatment
post-treatment
% Tethet CD45.1+CD45.2+
% Tetwt CD45.1+
Pre-T
Post-T
**
*
ns

Supplementary Figure 1

(A) In lethally-irradiated B6 CD45.1$^+$ recipient mice rescued with CD45.2$^+$ Tet2$^{wt}$ (n=4) or CD45.2$^+$ Tet2$^{het}$ bone marrow (n=5), the percent of CD45.2$^+$, donor-derived, tumor-infiltrating myeloid (CD11b$^+$), T cells (CD3$^+$), B cells (B220$^+$), or NK cells (NK1.1$^+$) were measured in MC38 tumors. There were no significant differences in the frequency of donor-derived cells between Tet2$^{wt}$- and Tet2$^{het}$–rescued mice by unpaired, two-tailed Student's T test.
(B) In lethally-irradiated B6 CD45.1$^+$ recipient mice rescued with CD45.2$^+$ Tet2$^{wt}$ (n=4) or CD45.2$^+$ Tet2$^{het}$ bone marrow (n=5), the distribution of TIL subtypes (both CD45.1$^+$ and CD45.2$^+$) in MC38 tumors expressed as a fraction of all CD45$^+$ TILs. There were no significant differences between between Tet2$^{wt}$- and Tet2$^{het}$–rescued mice by unpaired, two-tailed Student's T test.
(C) MC38 tumor growth in CD45.1 recipient mice rescued from lethal irradiation with CD45.2$^+$ Tet2$^{wt}$ (n=4) or Tet2$^{het}$ bone marrow (n=5).
(D) Mean tumor volume in mice with Tet2$^{wt}$ (black), Tet2$^{het}$ (red), or Tet2$^{KO}$ (blue) hematopoiesis treated with PD-1 ICB wt n=31; Tet2$^{het}$ n=32; Tet2$^{KO}$ n=14 over 3 experiments. * denotes day 24 $p < 0.05$ by two-tailed Mann-Whitney Test.
(E) Mean tumor volume in mice with Tet2$^{wt}$ (black), Tet2$^{het}$ (red), or Tet2$^{KO}$ (blue) hematopoiesis treated with isotype control antibodies. Tet2$^{wt}$ n=13; Tet2$^{het}$ n=13 iso; Tet2$^{KO}$ n=14 over 2 experiments. * denotes day 24 $p < 0.05$ by two-tailed Mann-Whitney Test.
(F) Tumor volume over time for each rescued genotype of isotype control treated mice, with the fraction of responders with endpoint tumor volume < 2 mm$^3$. p value from Fisher's Exact Test versus Tet2$^{wt}$.
(G) Flow cytometry gating strategy for myeloid leukocyte quantification (from live cells).
(H) Flow cytometry gating strategy for T and B cell quantification (from live cells).
(I) Flow cytometry gating strategy for hematopoietic stem and progenitor quantification (from live cells).
(J) CD45$^+$CD45.2$^+$ Tet2$^{het}$ and CD45.1$^+$ Tet2$^{wt}$ donor-derived bone marrow cells (schematic Figure 1D) in chimeric mouse bone marrow pre-treatment (light gray, n=4) and post-PD-1 ICB treatment (dark gray, n=5) as a percentage of all cells. Lineage negative, Sca1$^+$, c-Kit$^+$ (LSK), Multipotent progenitor (MPP) 2: LSK CD48+CD150+CD135-, , MPP3: LSK CD48+CD150-CD135-, MPP4: LSK CD48+CD150-CD135+, and common lymphoid progenitor (CLP): Lin-Sca-1$^{low}$c-Kit$^{low}$ CD127+CD135+ pre- and post-treatment frequency was compared for each genotype using unpaired, two-tailed Student's T test. ns, *, ** represent $p > 0.05$, $< 0.05$, and $< 0.01$, respectively.

Supplementary Figure 2

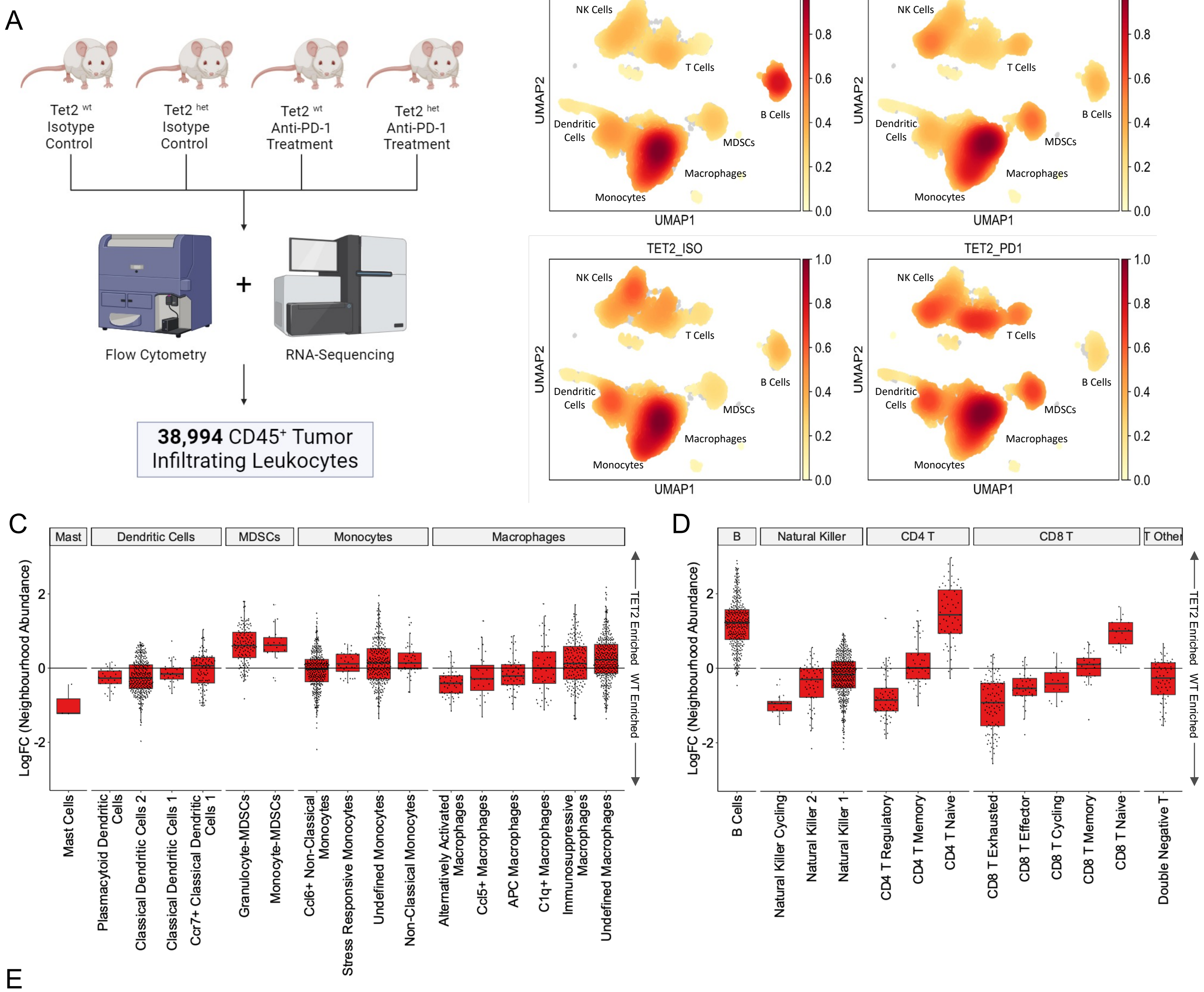


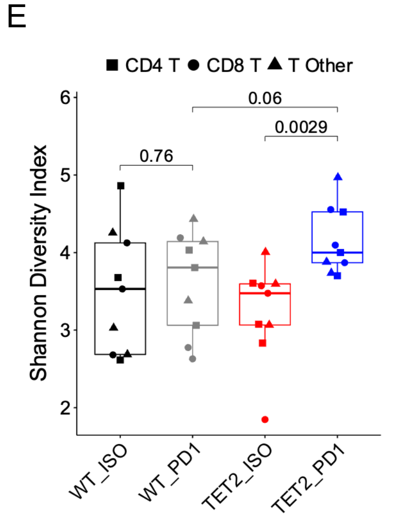


Supplementary Figure 2

(A) Experimental design for scRNA-seq. CD45+ tumor-infiltrating leukocytes from anti-PD-1 blockade treated (PD1, n=5 per genotype) or isotype control treated (ISO, n=3 per genotype) MC38 tumors in mice with $Tet2^{wt}$ or $Tet2^{het}$ hematopoiesis were isolated by FACS and underwent 10X 3' scRNA-seq and TCR VDJ analysis.

(B) UMAP projection displaying cell cluster density in TILs from $Tet2^{wt}$ (WT) and $Tet2^{het}$ TET2) mice treated with isotype control (ISO) or anti-PD-1 blockade (PD1).

(C) Lymphoid and myeloid (D) TIL subtype abundance displayed as log fold change between $Tet2^{wt}$ and $Tet2^{het}$ TILs from Milo neighborhood analysis of isotype control treated tumors. Fold change is in $Tet2^{het}$ mouse tumors relative to $Tet2^{wt}$ control tumors.

(E) Shannon Diversity Index from TCR VDJ sequencing analysis of $Tet2^{wt}$ and $Tet2^{het}$ (TET2) T Cells from isotype control (ISO) or anti-PD-1 (PD1) treated MC38 tumors. p values from two-tailed unpaired t-test.

Supplementary Figure 3

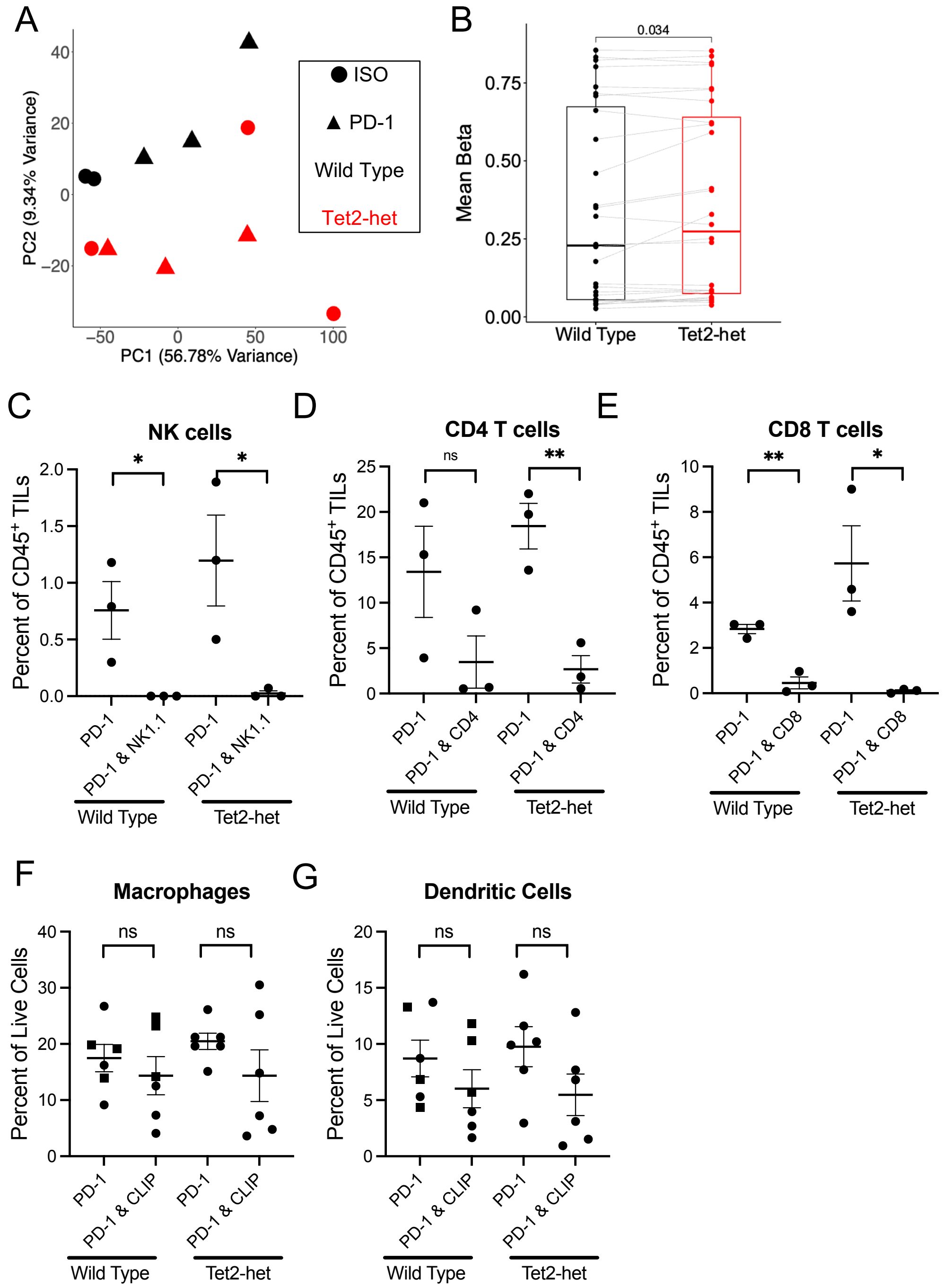


Supplementary Figure 3

(A) Principal component (PC) analysis of Illumina MM285 methylation array beta values in $Tet2^{wt}$ (black) or $Tet2^{het}$ (red) $F4/80^{+}$ cells from isotype control (ISO, circles) and anti-PD-1 (PD-1, triangles) treated MC38 tumors.
(B) Average MM285 methylation values (beta) of probes in the promoters of genes defining the scRNA-seq immunosuppressive macrophage cluster (Figure 2A) compared by two-tailed unpaired t-test between PD-1 treated $Tet2^{wt}$ (n=3) and $Tet2^{het}$ (n=3) $F4/80^{+}$ MC38 TILs.
(C-G) TIL subtype abundance in MC38 endpoint tumor dissociations from mice treated with anti-PD-1 blockade alone (PD-1), or anti-PD-1 blockade plus NK1.1 targeted antibody (C), CD4 depleting antibody (D), CD8 depleting antibody (E), or clodronate liposomes (CLIP) ($Cd45^{+}$,$F4/80^{+}$ macrophages F), ($Cd45^{+}$,$Cd11c^{+}$ dendritic cells G). *,**,ns denotes p value <0.05, 0.01, not significant, respectively, from 2-tailed, unpaired t-test.

### Supplementary Figure 4

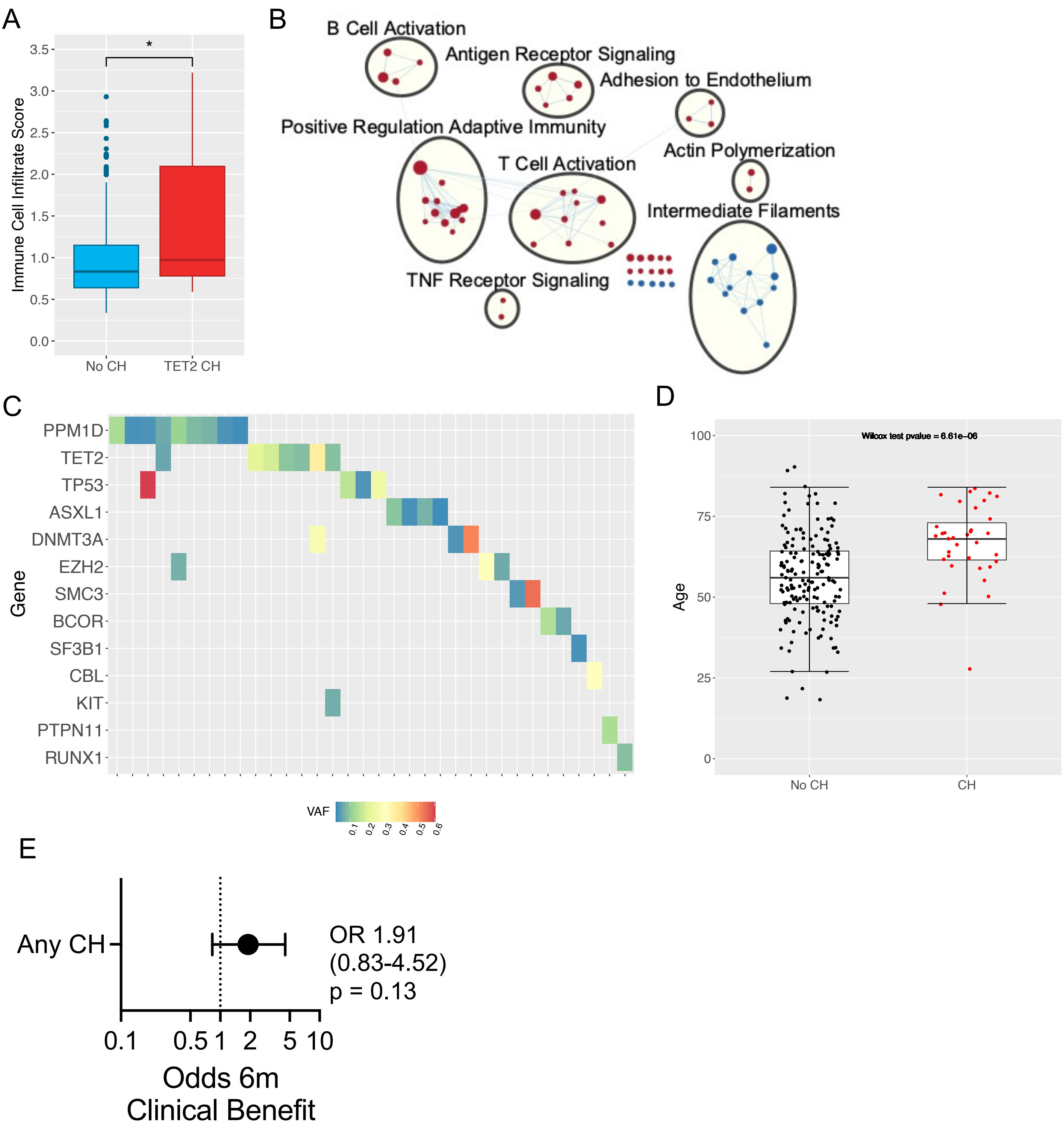

Supplementary Figure 4

(A) CIBERSORT$_x$ immune infiltrate scores compared between bulk-RNA-sequencing of Melanoma tumours from TCGA patients without CH (No CH) and with TET2-CH. * denotes p value < 0.05 from Mann-Whitney U test.

(B) Cytoscape Enrichment Map with AutoAnnotate annotation of genesets enriched in TET2-CH positive melanoma tumors bulk RNA-sequencing shown in red circles with genesets enriched in CH negative shown in blue, and lines representing connections between enriched genesets.

(C) CH driver gene mutations identified in PBMC or whole blood exome sequencing data from 203 ICB-treated metastatic melanoma patients with mutation variant allele frequency (VAF) shown.

(D) Average age of ICB-treated metastatic melanoma patients without (no CH) and with CH identified (CH) in PBMC or whole blood exome sequencing. Wilcoxon test p value shown comparing mean age in each group.

(E) Odds ratio of 6 month clinical benefit in metastatic melanoma patients exposed to CH versus no CH with 95% confidence interval, using Firth's penalized logistic regression with age, sex, study, and immune Checkpoint as covariates.
